# Estimating and testing the microbial causal mediation effect with high-dimensional and compositional microbiome data

**DOI:** 10.1101/692152

**Authors:** Chan Wang, Jiyuan Hu, Martin J. Blaser, Huilin Li

**Affiliations:** Division of Biostatistics, Department of Population Health, New York, University School of Medicine, New York, NY 10016, U.S.A.; Center for Advanced Biotechnology and Medicine, Rutgers University, Piscataway, NJ 08854-8021, U.S.A.

**Keywords:** Causal mediation analysis, Composition, Dirichlet regression, High-dimension, Hypothesis testing, Microbiome

## Abstract

**Motivation:** Recent microbiome association studies have revealed important associations between microbiome and disease/health status. Such findings encourage scientists to dive deeper to uncover the causal role of microbiome in the underlying biological mechanism, and have led to applying statistical models to quantify causal microbiome effects and to identify the specific microbial agents. However, there are no existing causal mediation methods specifically designed to handle high dimensional and compositional microbiome data.

**Results:** We propose a rigorous Sparse Microbial Causal Mediation Model (SparseMCMM) specifically designed for the high dimensional and compositional microbiome data in a typical three-factor (treatment, microbiome and outcome) causal study design. In particular, linear log-contrast regression model and Dirichlet regression model are proposed to estimate the causal direct effect of treatment and the causal mediation effects of microbiome at both the community and individual taxon levels. Regularization techniques are used to perform the variable selection in the proposed model framework to identify signature causal microbes. Two hypothesis tests on the overall mediation effect are proposed and their statistical significance is estimated by permutation procedures. Extensive simulated scenarios show that SparseMCMM has excellent performance in estimation and hypothesis testing. Finally, we showcase the utility of the proposed SparseMCMM method in a study which the murine microbiome has been manipulated by providing a clear and sensible causal path among antibiotic treatment, microbiome composition and mouse weight.

## 1 Introduction

Microbiome research is producing exciting results, with many studies linking specific microbes to particular diseases (Gilbert *et al.*, 2018; Ni *et al.*, 2017; Zheng *et al.*, 2016), physiological properties and environmental parameters (Albenberg and Wu, 2014; Stein *et al.*, 2016; Zeevi *et al.*, 2015). However, knowing the correlation or association between microbiome and another trait (Hu *et al.*, 2018; Koh *et al.*, 2017) is no longer a sufficient research goal, since now the scientific frontier is to understand the causal role of microbiota in the underlying biological mechanism (Fischbach, 2018). For example, early life exposure to antibiotics has been reported to affect human metabolism and cause weight gain and/or immunological properties by altering the gut microbiota (Livanos *et al.*, 2016; Mahana *et al.*, 2016; Schulfer *et al.*, 2018, 2019). To confirm such causal relationships, researchers conducted experiments randomizing groups of newborn mice control and antibiotic groups, and collecting longitudinal microbiome, weight, and immunological measurements through the study period. Within each experiment, researchers sought to understand how the change in microbiome due to the antibiotic exposure caused the change in mouse phenotype. If the altered microbiome played a causal role, which specific microbes were the culprits? To answer these questions, a rigorous causal mediation analytic framework is needed.

In microbiome causal research, there are many possible factors can potentially confound the microbiome effect, such as age, gender and diet (Knight *et al.*, 2018). In order to control or eliminate the possible confounding effects, randomized experiments are usually used to investigate the specific effects of a course of treatment on the microbiome and diseases (Knight *et al.*, 2018). In this paper, we introduce our proposed microbiome causal analytical framework within randomized experimental designs and target understanding the causal pathway among three factors: treatment (T), microbiome (**M**) and outcome (Y) (**Fig. 1**). Here, microbiome are hypothesized as mediators on the pathway from treatment to outcome. It is important to understand how the causal effect of the treatment on the outcome can be divided into the causal direct effect and the causal indirect effect, acting through the mediator (the latter is also called the causal mediation effect in this paper).

**Figure 1:**
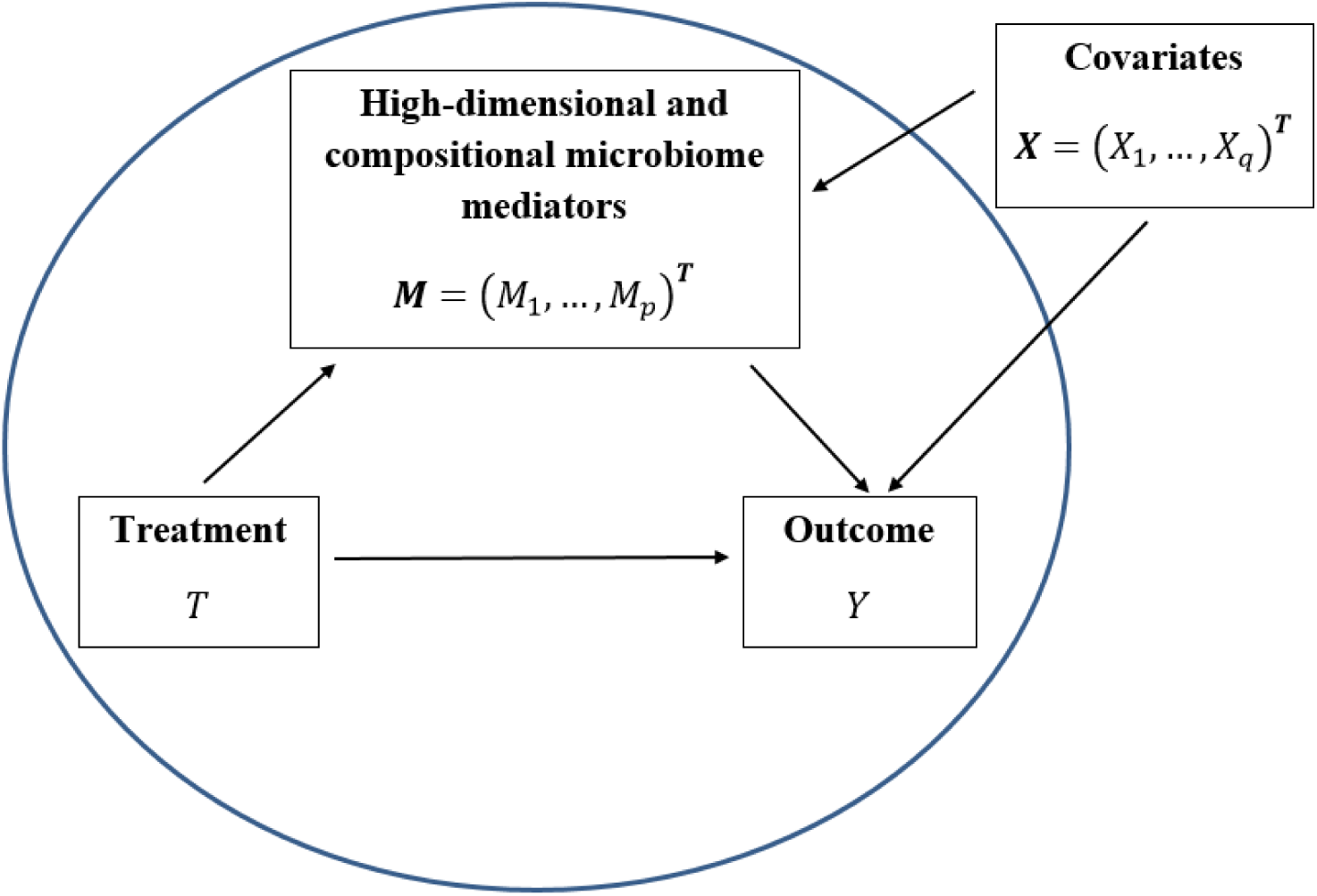
Relations between treatment *T*, covariates ***X***, microbiome composition (mediators) ***M*** and outcome *Y*. We aim to investigate the effect of treatment directly on the outcome (termed as the direct effect) and the effect of treatment through microbiome composition as mediators (termed the mediation effect), while adjusting for covariates.

Donald Rubin and Paul Holland developed the potential outcome idea (Neyman, 1923) and established a formal mathematical causal framework for both observational and randomized experimental studies. (Holland, 1986; Rubin, 1974, 2005). They defined causation as a hypothetical value through the counterfactual statement: a score difference between the observed outcome of one subject under one treatment condition and the potential outcome if he/she would be under the alternative treatment condition. Later, VanderWeele and his colleagues expanded this framework to the mediation analysis to define causal direct effect and causal mediation effect under the sufficient causal assumptions and allow for the presence of exposure-mediator interactions (Valeri and VanderWeele, 2013; VanderWeele and Vansteelandt, 2014; VanderWeele, 2013, 2014; VanderWeele and Vansteelandt, 2009, 2010). In our motivating example and other microbiome research, it is possible that the treatment can affect microbiome’s mediating effect on the outcome, so we follow VanderWeele’s counterfactual mediation framework and incorporate the interaction of treatment and micrbiome on the outcome into the proposed method.

Recently, with the advent of high-throughput biomedical data, a few causal mediation models have become available to handle high-dimensional mediators by using linear structural equation modeling (LSEM). These methods adopted two ways to reduce the dimensionality of the mediators. One way is through regularization or penalization. For example, by assuming that there is no correlation among mediators, Zhang *et al.* (2016) proposed a joint significance test based on sure independent screening (Fan and Lv, 2008) and minimax concave penalty techniques (Zhang *et al.*, 2010) to evaluate the casual mediating effect of DNA methylation markers. The other way is to transform the correlated high-dimensional mediators into a series of causal mediation models with single continuous mediator. For example, Huang and Pan (2016) used spectral decomposition to transform high-dimensional gene expression mediators into low-dimensional and uncorrelated ones. Chén *et al.* (2017) transformed high-dimensional imaging mediators into orthogonal components and ranked them based on their contributions to the LSEM likelihood.

Unfortunately, none of the methods cited above is designed for the high dimensional, sparse, and compositional microbiome data. Compositionality is the crux of the new challenges in causal studies involving the microbiome. Due to varying sequencing read counts across samples, normalization needs to be employed to make the microbial counts comparable before downstream analyses. As a common normalization method, the sequence counts are scaled by the total number of reads or the total number of reads and lengths of reads together. This step converts the count data into the relative abundance (Knight *et al.*, 2018), which is compositional and has the simple unit-sum constraint. Log-ratio analysis (Aitchison, 1982) and Dirichlet regression (Hijazi and Jernigan, 2009) are two available methods to deal with relative abundance. Aitchison’s log-ratio methods work on the ratios of the components of the composition. Because those ratios are sensitive to low relative abundance and need additional handling for zero counts, the log-ratio methods have been criticized for its limited interpretability (Hijazi and Jernigan, 2009). As an alternative, Dirichlet regression has drawn attention by analyzing compositional data with multivariate statistical modeling, and has been shown to be useful for compositional data (Campbell and Mosimann, 1987a,b; Hijazi and Jernigan, 2009).

In the paper, we propose a sparse microbial causal mediation framework (SparseMCMM) to clarify the relationship among a binary treatment, a vector of compositional microbial mediators, and a continuous outcome. We use Dirichlet regression to model the relationship between treatment and microbiome composition (Hijazi and Jernigan, 2009), and linear log-contrast regression to model treatment effect, log transformed microbiome effects, and the effects of their interactions on the outcome (Aitchison and Bacon-shone, 1984; Lin *et al.*, 2014). The combination of those two regressions are then considered in the counterfactual mediation framework to clarify the causal mediation effect of microbiome on the outcome.

The remainder of this paper is organized as follows: In Section 2, we introduce the proposed SparseMCMM with high-dimensional and compositional mediators under the counterfactual framework and derive the corresponding mediation effect estimators. Then, we describe how to select and estimate non-zero parameters in the high-dimensional causal mediation regressions with regularizations. Moreover, we propose two hypothesis tests for the overall mediation effect and apply permutation procedures to access their statistical significance. In Section 3, we conduct extensive simulations to evaluate the performance of the proposed model on both estimation and testing. Subsequently, we implement SparseMCMM to investigate the extent to which the microbiome mediates the causal pathway from antibiotic usage to excess weight gain in a longitudinal murine study. We conclude with discussion of the relevant issues in Section 4.

## 2 Methods

### 2.1 Casual mediation model

#### 2.1.1 Notations and two basic regression models

Suppose there are *n* subjects, *p* taxa, and *q* covariates and subscripts *i, j, k*, indicate a subject, a taxon, and a covariate, respectively. For the *i*th subject, let *T*_*i*_ be the treatment status with *T*_*i*_ = 1 or 0 for the treatment group or the control group, ***M***_*i*_ = (*M*_*i*1_, …, *M*_*ip*_)^*T*^ be the microbiome relative abundance with the constraint 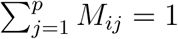, let ***X***_*i*_ = (*X*_*i*1_, …, *X*_*iq*_)^*T*^ represent the covariates such as age, sex, and weight, and let *Y*_*i*_ be the continuous outcome. We propose two regression models to model the causal mediation relationships among *T*, ***M***, ***X***, and *Y* as illustrated in **Fig. 1**. The first model depicts that the outcome *Y*_*i*_ is determined by the treatment, compositional mediators, interactions between the treatment and mediators, and covariates as:

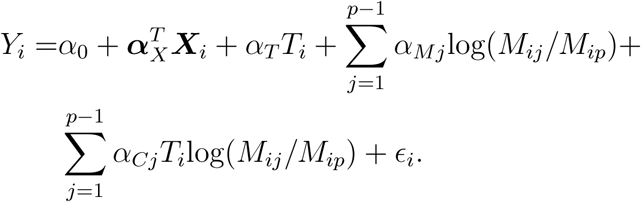

Note that with 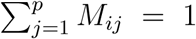, the relative abundances of *p* taxa are built into the model through the log-contrast strategy with the *p*th taxon as the reference using the log ratio transformation proposed by Aitchison and Bacon-shone (1984). By defining 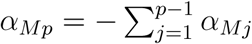 and 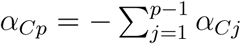 (Lin *et al.*, 2014), with algebraic equivalents, we can rewrite the above model in the matrix form,

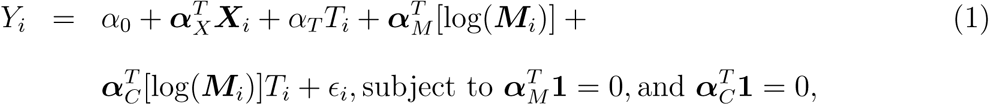

where *α*_0_ is the intercept, *α*_*T*_ is the coefficient of treatment, ***α***_*X*_ = (*α*_*X*1_, …, *α*_*Xq*_)^*T*^, ***α***_*M*_ = (*α*_*M*1_, …, *α*_*Mp*_)^*T*^, and ***α***_*C*_ = (*α*_*C*1_, *…*, *α*_*Cp*_)^*T*^ are the vectors of coefficients of covariates, microbial mediators, and interactions between treatment and mediators, respectively, and *ϵ*_*i*_ ∼ *N* (0, *σ*^2^) is the error term.

Secondly, we use the Dirichlet regression (Hijazi and Jernigan, 2009) to model the microbial relative abundance as a function of treatment and covariates. Specifically we assume that ***M***_*i*_|(*T*_*i*_, ***X***_*i*_) ∼ Dirichlet (*γ*_1_(*T*_*i*_, ***X***_*i*_), *…*, *γ*_*p*_(*T*_*i*_, ***X***_*i*_)), and their microbial relative means are linked with treatment and covariates (*T*_*i*_, ***X***_*i*_) in the generalized linear model fashion with a log link:

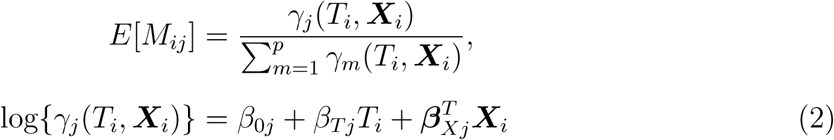

where *β*_0*j*_ is the intercept and *β*_*T j*_ and ***β***_*Xj*_ are the coefficients of treatment and covariates for the *j*th taxon, respectively. With this modeling, we quantify the treatment effect on each individual taxon which further allows the quantitation of component-wise (or taxon-wise) mediation effect and the overall (or aggregated) mediation effect of the microbiome community in the next subsection.

#### 2.1.2 Definition of direct and mediation effects in the counterfactual framework

With the above two models, we next determine the average causal direct effect of treatment and the average mediation effect of microbiome on the outcome under the counterfactual framework (VanderWeele and Vansteelandt, 2014; VanderWeele, 2016; VanderWeele and Vansteelandt, 2009). With counterfactual notation, DE refers to the expected difference of the outcome *Y* between the treatment *T* = 1 and *T* = 0 when the mediators ***M*** are set to the value they would have taken had *T* been set to 0, and is defined as:

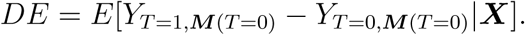

ME is the indirect effect of treatment on outcome through the compositional microbiome community and refers to the expected difference of the outcome *Y* between the mediators ***M*** (*T* = 1)|***X*** and ***M*** (*T* = 0)|***X*** when the treatment *T* = 1, where ***M*** (*T* = *t*)|***X*** represents the microbiome composition we would have observed had *T* been set to the value *t* given covariates ***X***. Mathematically, ME is defined as:

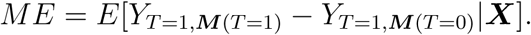

Under four sufficient identifiable assumptions (see Supplementary Materials, Section S1), DE and ME can be further expressed by the parameters in models (1)-(2), respectively, as follows:

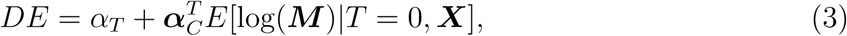

and

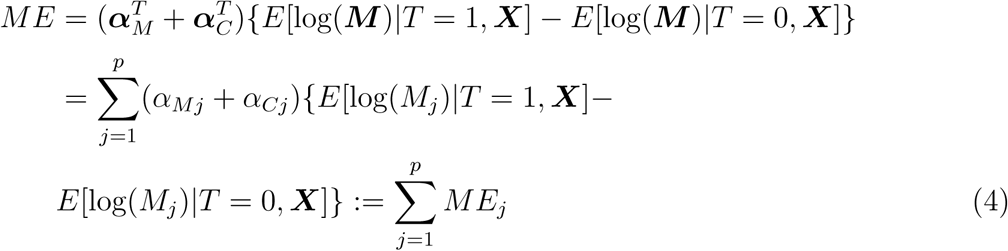

From equation (4), note that ME is the summation of the individual mediation effects from each taxon *ME*_*j*_. *ME*_*j*_ is the product of two parts: (*α*_*Mj*_ +*α*_*Cj*_) which represents the *j*th microbial effect consisting of the main effect and the interaction effect of this taxon and the treatment on the outcome; and *{E*[log(*M*_*j*_)|*T* = 1, ***X***] *- E*[log(*M*_*j*_)|*T* = 0, ***X***]*}* which represents the treatment effect on the *j*th taxon. Therefore *ME*_*j*_ exists only if when both the *j*th microbial effect on the outcome and the treatment effect on the *j*th taxon are not zero. For the expectation part, we have 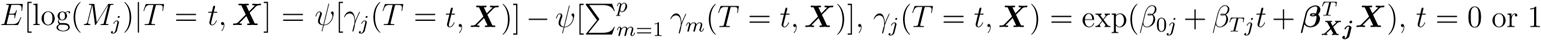, and 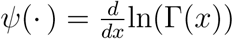 is the digamma function.

The total effect of the treatment on the outcome is therefore the summation of DE and ME:

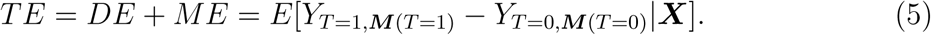

The detailed derivations of DE, ME and TE are provided in the Supplementary Materials, Section S2.

In summary, given covariates ***X***, SparseMCMM is able to decompose the treatment effect on the outcome into the direct effect of treatment and the mediation effects through the microbiome. It elucidates the mediation role of the microbiome through rigorous statistical modeling, and quantifies the mediation effects for the overall microbiome community and for each specific taxon respectively.

### 2.2 Parameter estimation

It is challenging to estimate all parameters from the joint log-likelihood function based on models (1)-(2) due to the nonlinearity and constraints. As an alternative approach, we first estimate the regression parameters in models (1)-(2), separately and then include them into equations (3)-(5) to obtain the estimated DE, ME (*ME*_*j*_ for the individual taxon) and TE, respectively. A similar two-step approach has been used in the genetics field and its computing efficiency has been recognized (Huang and Pan, 2016; Zhang *et al.*, 2016).

Another challenge of estimation in the microbiome setting is the high dimensional mediators. Model (1) has ∼ 2*p* parameters when it considers both main effect and interaction effect and model (2) has ∼ (*q* + 2)*p* parameters. Both models have far greater number of parameters than the sample size *n*. To deal with the hign-dimensional mediators, we propose regularization techniques to simultaneously identify the key taxa with the primary mediation effect and parameter estimation. Specifically, in model (1), we propose a penalized least squares criterion to penalize the main effects and interaction effects of mediators, which has optimal biological explanations; in model (2), we utilize the *L*_1_ norm penalty to select taxa which are altered by the treatment. In the following section, we introduce the estimation procedures for models (1) and (2) sequentially.

#### 2.2.1 Parameter estimation for the linear log-contrast regression

Denote the parameters of regression coefficients in model (1) by 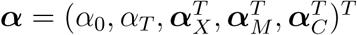. We use the least squares method (Friedman *et al.*, 2001) to estimate ***α***. Given the observed data (*T*_*i*_, ***X***_*i*_, ***M***_*i*_, *Y*_*i*_), the sum of squared residuals (SSR) which measures the discrepancy between the observed and predicted outcome is:

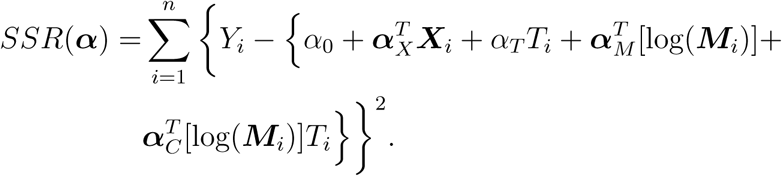

It is a well-established variable selection practice in high dimensional linear regression with interaction that the interaction effect exists only if the corresponding main effects are included in the model, which is termed the heredity condition or hierarchy structure (Peixoto, 1987; Radchenko and James, 2010). To apply this constraint, we add two penalties to the SSR and solve the following convex least squares optimization problem:

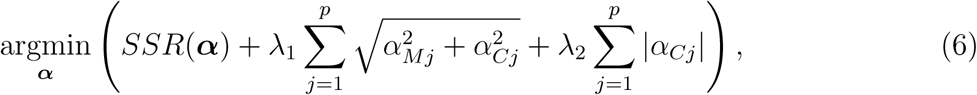

with the constraints 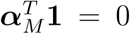 and 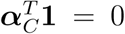. *λ*_1_(*≥* 0) and *λ*_2_(≥ 0) are two tuning parameters. As discussed by Radchenko and James (2010), these penalty functions have desirable properties in both theory and performance in addressing the heredity condition. Specifically, the first penalty, similar to the group-lasso penalty, ensures that the main effect *α*_*Mj*_ and the interaction effect *α*_*Cj*_ of the *j*th taxon shrinks on the same scale. The second *L*_1_ norm penalty only works on the interaction terms. In combination, they guarantee that the main effect *α*_*Mj*_ from the first penalty could only be shrunk to 0 when the corresponding interaction effect *α*_*Cj*_ from the second penalty also is shrunk to 0. When *α*_*Cj*_ = 0, the first penalty is reduced to a lasso penalty: 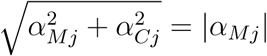.

We utilize the sequential quadratic programming (SQP) method, a popular method for solving constrained nonlinear optimization problems, with R package nloptr (Kraft, 1988; Ypma, 2014) to optimize equation (6) and obtain the estimate 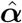. Particularly, SQP seeks a numerical solution by solving a sequence of quadratic sub-problems, each of which optimizes a quadratic objective function subject to the linear constraints. Tuning parameters *λ*_1_ and *λ*_2_ are determined by Bayesian information criterion (BIC) (Chen and Chen, 2008).

#### 2.2.2 Parameter estimation for the Dirichlet regression

Denote the parameters of regression coefficients in model (2) by 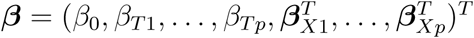. Given the observed data (*T*_*i*_, ***X***_*i*_, ***M***_*i*_), the log-likelihood function for *n* observations is given by

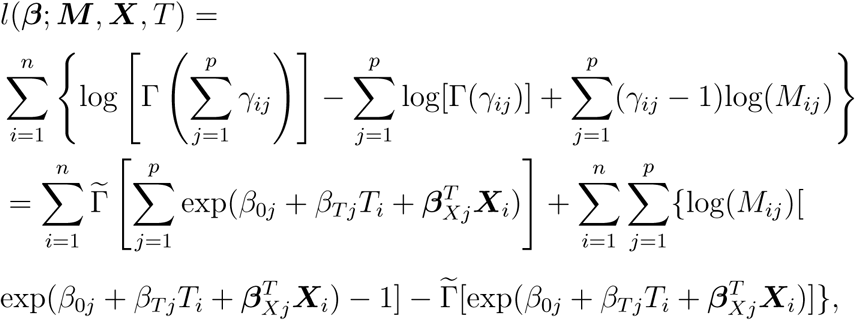

where *γ*_*ij*_ = *γ*_*j*_(*T*_*i*_, ***X***_*i*_) as defined in model (2), and 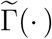 is the log gamma function. In order to select the taxa whose relative abundances are altered by treatment, we minimize the following penalized version of the log-likelihood with *L*_1_ penalty:

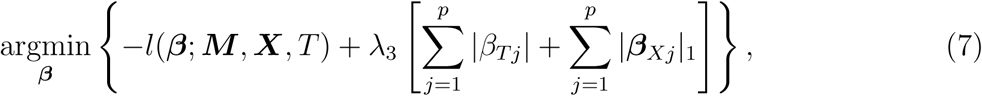

where *λ*_3_ ≥ 0 is the tuning parameter and is determined by BIC as those in model (6). The Newton-Raphson algorithm in nloptr R package (Bonnans *et al.*, 2006) is used to find the numerical estimate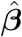.

Inserting the estimates 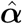 and 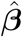 into equations (3)-(5), we can obtain the estimates 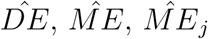 and 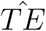 respectively.

### 2.3 Hypothesis tests for mediation effects

We propose two tests to examine whether the microbiome has any mediation effect on the outcome or not, at the community and taxon levels, denoted as OME and CME, respectively, in equation (4).

Since the null hypothesis of no overall mediation effect at the community level can be expressed *H*_0_: *ME* = 0, the first test is defined as

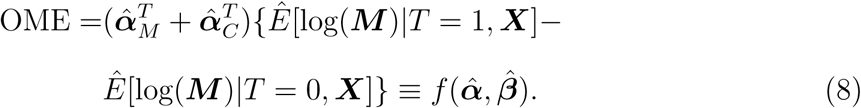

From equation (4), we can see that the overall mediation effect of the microbiome community is defined as the summation of all component-wise *ME*_*j*_s and it can be counteracted when both positive and negative component-wise *ME*_*j*_s are present. Thus, in the second test, we consider the following null hypothesis which tests whether at least one component-wise *ME*_*j*_ is significantly non-zero:

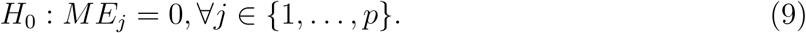

To tackle this problem with the high dimensionality of the mediators, we propose an equivalent null hypothesis 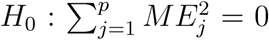, as indicated by Huang and Pan (2016). Then CME test is formulated as

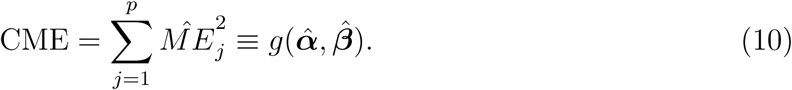

It is not trivial to derive the asymptotic distributions of the test statistics OME and CME. As an alternative, we apply the following permutation procedure to estimate p-values (Boca *et al.*, 2013; Taylor and MacKinnon, 2012; Zhang *et al.*, 2018).

First, we randomly shuffle treatment *T* and outcome *Y* separately to obtain the permuted treatment *T* ^(*b*)^ and outcome *Y* ^(*b*)^, *b* = 1, *…*, *B*. Second, we get estimates 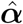 and 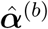 from model (1) based on data (*T*, ***X, M***, *Y*) and (*T*, ***X, M***, *Y* ^(*b*)^) respectively, and 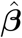 and 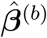 from model (2) based on data (*T*, ***X, M***) and (*T* ^(*b*)^, ***X, M***) respectively. Third, we calculate three sets of permuted statistics: 1.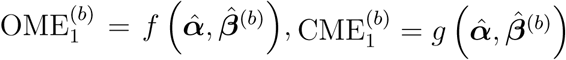; 2.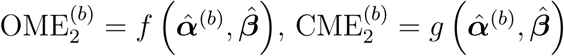; and 3. 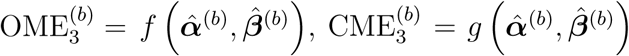 to cover three different types of null hypotheses: 1. *T* ⇏ **M** ⇒*Y*; 2. *T* ⇒ **M** ⇏ *Y*; and 3. *T* ⇏ **M** ⇏ *Y* in testing the mediation effect, respectively. And the final test statistics for the *b*^th^ permutation are defined as

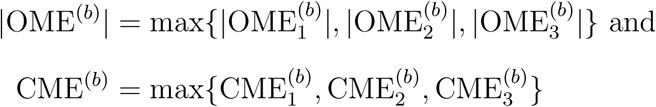

respectively. Therefore, the testing p-values for OME and CME are

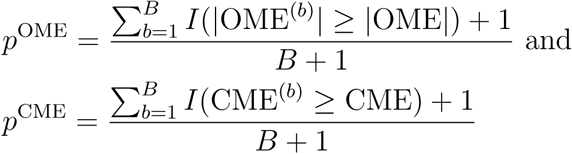

respectively, where *I*(·) is the indicator function.

## 3 Results

### 3.1 Simulation studies

We conducted extensive simulation studies to evaluate: 1) the estimation performance of SparseMCMM in terms of bias and mean squared error (MSE) for DE and ME, respectively; and 2) the testing performance of SparseMCMM in terms of the empirical type I error rate and power, compared with tests Delta.T and tau.T proposed in Huang and Pan (2016) and HIMA proposed in Zhang *et al.* (2016), representing regularization-based and transformation-based tests to handle high-dimensional mediators respectively. Since these competing tests are designed to deal with continuous mediators, rather than the compositional microbiome data, we log transformed the relative abundances to make them more normal. Detailed introduction for these three competing tests is provided in the Supplementary Materials, Section S3.

#### 3.1.1 Simulation design

We designed our simulation settings based on the experimental design of the murine microbiome study we analyzed as the real data example in Section 3.2 (Schulfer *et al.*, 2019). To make the simulation simple and focused, we did not consider any covariates in the simulation, although our SparseMCMM package has the full capacity to handle any covariate adjustment. We generated the simulation data in three steps: 1) generate the treatment *T*; 2) generate the microbiome/mediators ***M*** based on treatment *T*; and 3) generate the outcome *Y* based on (*T*, ***M***). The detailed simulation design is provided below.

##### Generate the treatment T

for *n* total subjects, we randomly assigned 50% to treatment group (*T* = 1) and the others to control group (*T* = 0). *n* = 50, 100, 300, and 500 for evaluating estimation and *n* = 100 for evaluating testing.

##### Generate the microbiome M

we simulated microbiome data based on the Dirichlet distribution reflecting the real microbial composition in our murine example. The mean relative abundances of *p* taxa for the treatment and control groups follow the simplified model (2), as below:

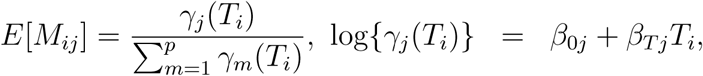

where ***β***_0_ = (*β*_01_, *…*, *β*_0*p*_)^*T*^ represents the log-transformed baseline relative abundances for *p* taxa and they were set as the corresponding estimates from the male control mice at day 28 (using R package dirmult; Tvedebrink (2009)). The real data include 149 genera and we randomly divided them into *p* taxa. The specific values of ***β***_0_ with *p* = 10, 25 and 50 used in the simulations are listed in **Table S1**. For the treatment group, we randomly chose *p*_*r*_ out of *p* taxa as the causal ones (*p*_*r*_ *< p*) and denote the set of their indices as ^ and their treatment coefficients as 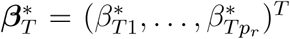. For those non-causal taxa, *β*_*T j*_ = 0, *j* ∉ ^. We considered various sets of 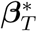 to represent different simulation settings. With the specific ***β***_0_ and ***β***_*T*_ values, microbiome composition data ***M***_*i*_ could be generated from the Dirichlet distribution.

##### Generate the outcome Y

the outcome *Y* was generated based on model (1) with the simulated treatment *T*, and microbiome composition ***M***. We set the regression coefficients of intercept, treatment and non-causal taxa as *α*_0_ = 0, *α*_*T*_ = 1 and *α*_*Mj*_ = *α*_*Cj*_ = 0, *j* ∉ ^, respectively. For *p*_*r*_ causal taxa, we denoted their regression coefficients for the main effect and interaction effect by 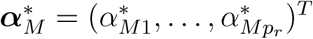 and 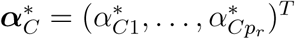, respectively, and their values were set differently to represent various simulation scenarios.

##### Specify 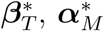 and 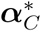

First, the number of causal taxa was set as *p*_*r*_ = 2, 3 and 5 for *p* = 10, 25 and 50 respectively.

1. To evaluate the performance of the DE and ME estimators, parameter values (**Table S2**) were set up as the corresponding estimates of male mice at day 28 in the real data analysis, so that the true TE and ME were around 1.8 and 0.6 respectively (**Table S3**). Finally, 1,000 independent replications were conducted to calculate bias and MSE for the DE and ME estimators. The corresponding computational time is given in **Table S4**.
2. To evaluate the performance of OME and CME tests, we considered two simulation scenarios: 1) all individual *ME*_*j*_s of the causal taxa are positive; and 2) both positive and negative *ME*_*j*_s are present. Further, four strength levels of ME: null, small, medium, and large were considered in each scenario. Note that null ME was used to evaluate the empirical type I error rate and the other three strength were used to evaluate the empirical power. Please see the parameter setting for 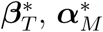 and 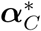 in **Table S5**. To ease our computation, we restricted this part of simulation to the sample size of 100. The p-values were estimated based on 500 permutations (**Table S6** reports the computational time), and then the empirical type I error rate and power were calculated by the proportion of p-values less than the given significance level (usually 0.05) with 1,000 independent replications.

#### 3.1.2 Estimations of causal direct effect and mediation effect

**Table 1** shows the bias and MSE for the DE and ME estimators respectively. First, as the dimension of the mediators *p* increases from 10 to 50, both bias and MSE of DE and ME estimators increase. In contrast, with each fixed *p*, as the sample size *n* goes up, the bias and MSE go down. When *n* = 500, the bias of both DE and ME estimators approach zero. This indicates that the DE and ME estimators are approximately unbiased. The proposed method, SparseMCMM, therefore has good performance in direct effect and mediation effect estimation.

**Table 1:** Bias and MSE of the proposed causal direct effect (DE) and mediation effect (ME) estimators for various sample sizes and dimensions of the compositional mediators.

| $p$ | $n$ | DE | | ME | |
| --- | --- | --- | --- | --- | --- |
|  |  | Bias | MSE | Bias | MSE |
| 10 | 50 | 0.0016 | 0.0093 | -0.0047 | 0.0091 |
|  | 100 | -0.0025 | 0.0043 | 0.0019 | 0.0047 |
|  | 300 | -0.0025 | 0.0013 | -0.0013 | 0.0017 |
|  | 500 | -0.0016 | 0.0009 | 0.00021 | 0.0011 |
| 25 | 50 | 0.062 | 0.092 | 0.020 | 0.35 |
|  | 100 | 0.026 | 0.053 | 0.019 | 0.22 |
|  | 300 | 0.021 | 0.051 | -0.010 | 0.16 |
|  | 500 | 0.012 | 0.031 | -0.0075 | 0.090 |
| 50 | 50 | 0.21 | 1.70 | -0.43 | 1.00 |
|  | 100 | 0.22 | 0.23 | -0.081 | 0.30 |
|  | 300 | 0.038 | 0.048 | -0.026 | 0.13 |
|  | 500 | 0.017 | 0.037 | -0.0039 | 0.12 |

**Fig. S2** exhibits the boxplot of 1,000 estimated component-wise *ME*_*j*_s for *j* = 1, *…*, *p*, with *p* = 10, 25, 50 and *n* = 50, 100, 300, 500 respectively. The bias and MSE of component-wise *ME*_*j*_s clearly decrease, as the sample size increases. The figures show that for the non-causal taxa, their *ME*_*j*_ estimates all are around zeros, while for the causal taxa, their *ME*_*j*_ estimates all stand out and are away from zero except for taxon 22 when *p* = 50 (its true *ME*_22_=0.002). Overall, SparseMCMM presents good performance in both causal taxa selection and its estimations.

#### 3.1.3 Power and type I error rate

**Table S7** reports the empirical type I error rates of OME, CME, Delta.T, tau.T, and HIMA. They all are below the nominal significance levels 5%. OME and CME have conservative type I errors, which is consistent with the results in Boca *et al.* (2013) and Zhang *et al.* (2018). Delta.T and tau.T have relatively more conservative type I error rates than OME and CME do, which agrees with their conservative performance in the power section.

**Fig. 2** (*p* = 10, 50) and **Fig. S3** (*p* = 25) present the power estimations of OME, CME, Delta.T, tau.T, and HIMA for the same effect direction (scenario 1) and mixed effect directions (scenario 2) with small, medium and large overall MEs. Compared to Delta.T, tau.T, and HIMA, the proposed tests OME and CME have superior performances in both scenarios with *p* = 10, 25, and 50, and they exhibit an increasing power trend as the overall ME increases. For the comparison between OME and CME, as expected, OME gains more power than CME in scenario 1, when the component-wise *ME*_*j*_s for the causal taxa all are positive, but it is the reverse for scenario 2 when the directions of the individual causal effects are mixed. Among those three competing tests, HIMA has the best performance in both scenarios. However, none has comparable performance to OME or CME. This implies that one needs to be cautious when directly applying the existing high dimensional mediation methods, which are not designed for taking care of the unit sum constraint, to the compositional microbiome data.

**Figure 2:**
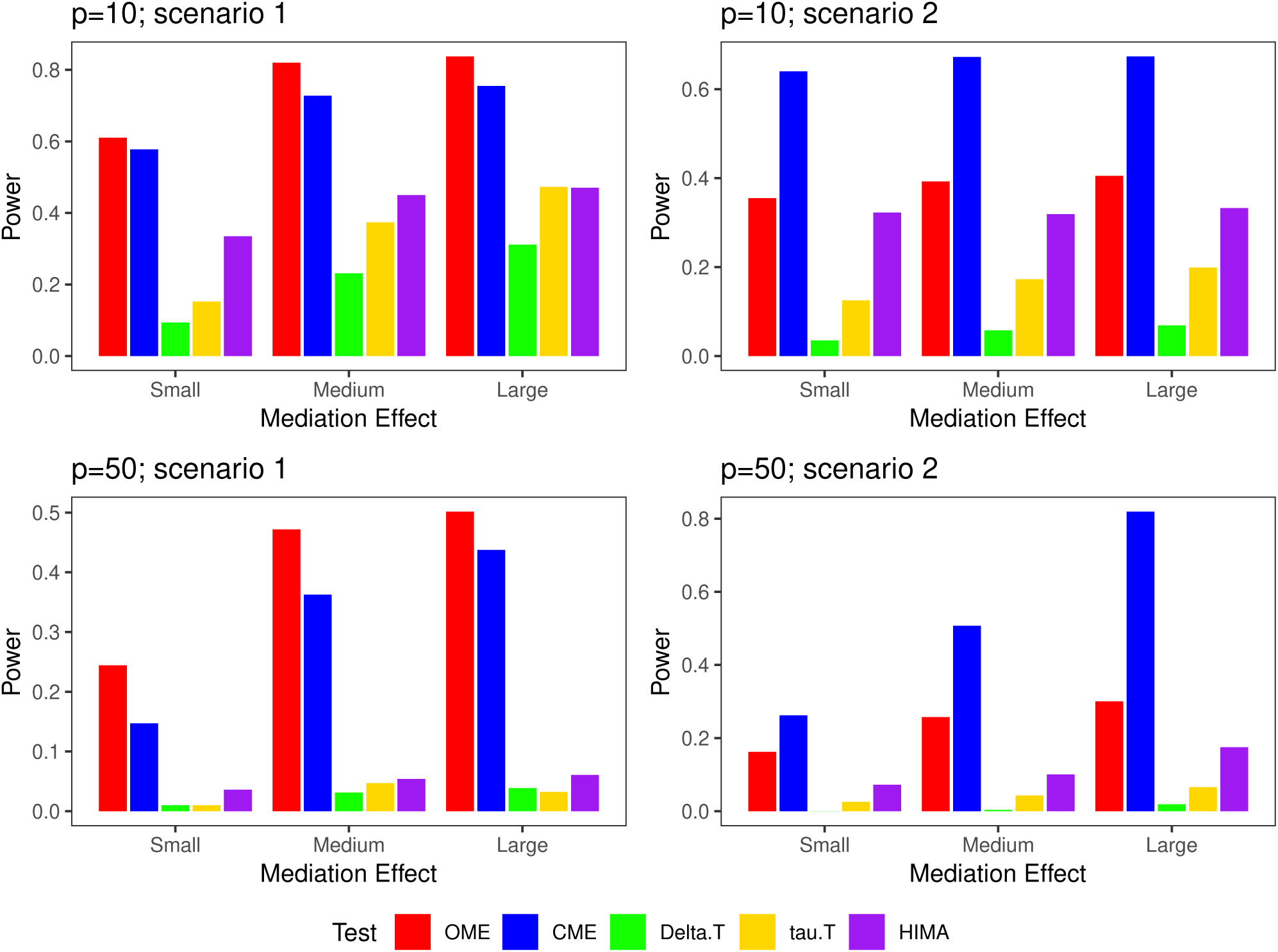
Empirical power for testing mediation effect with *p* = 10 and 50 in scenarios 1-2 (significance level=5%). Note that the magnitudes of mediation effect are not comparable across different *p*s and different scenarios. The detailed setting is given in **Table S5.**

### 3.2 Real data analysis

Schulfer *et al.* (2019) conducted a murine microbiome experiment to explore whether STAT (sub-therapeutic antibiotic treatment) would alter gut microbiome composition and whether this shift would change the body weight gain later in life. Since this study adopts the typical treatment-microbiome (mediator)-outcome design, we use SparseMCMM to re-examine the mediating role that the gut microbiome played in body weight gain. To be specific, we first use OME test to determine whether the overall mediation effect of microbiome is significant, and then use CME test to determine whether at least one individual taxon have significant mediation effects on the body weight gain. Subsequently, SparseMCMM gives the overall ME and individual *ME*_*j*_ estimates for microbiome.

In this study, DNAs were extracted from fecal samples using the 96-well MO BIO PowerSoil DNA Isolation Kit by targeting the V4 region of the bacterial 16S rRNA gene, as described in Caporaso *et al.* (2010). Samples with less than 1800 reads were excluded from the analysis. The OTU table for 21 female (12 STAT and 9 controls) and 37 male (24 STAT and 13 controls) mice was constructed using the QIIME pipeline (Caporaso *et al.*, 2010) at day 21 and 28. Originally there were 149 genera. After filtering those genera that appeared in *<* 10% of mice and with mean proportions *<* 10^*-*4^ at each time point separately, there were 38 and 37 genera retained at days 21 and 28 respectively. The observed body weight (in grams) prior to sacrifice, i.e., at day 145 for the female and at day 116 for the male mice was regarded as the outcome. No additional covariates were included in the model, assuming that all potential confounders had been well-controlled in the randomized experiment.

**Fig. S4** illustrates the distribution of the body weight prior to sacrifice in the control and STAT groups, respectively, for female (left panel) and male mice (right panel). Male mice are known to be heavier than the female mice, and within each gender, the STAT group was heavier than the control group. Considering these different weight distributions between genders and the sexually dimorphic effect of STAT in mice, we explored the mediation effect of gut microbiome on the acceleration of weight gain for female and male mice separately. Since that Delta.T and tau.T tests proposed in Huang and Pan (2016) have no significant results (Table S8), and HIMA proposed in Zhang *et al.* (2016) gives testing results only at the genera level (Table S9), we only discuss the results for the proposed causal mediation method SparseMCMM next.

**Table 2** reports the testing results for OME and CME based on 500 permutations. For the females, OME test shows that the overall mediation effect of microbiome is significant (p-value=0.048*<* 0.05) at day 28, but not significant at day 21 (p-value=0.154). The estimated overall ME increases from 0.102 (3% of total causal effect on the weight gain) at day 21 to 0.634 (18.8%) at day 28. On the other hand, CME test is also significant at day 28 with p-value 0.04, which shows that there is at least one genus playing a mediation role on the weight gain at day 28. With SparseMCMM, we further identify four candidate genera, reported in **Table 3** with the point and 95% confidence interval (CI) estimates for their mediation effects. **Table 3** also reports the breakdown of mediation effect estimates. Column 2 indicates the microbiome effect on the weight, column 3 indicates the treatment effect on the microbiome, and the multiplication of those two columns equals the mediation effect estimate for each genus in column 4. Among the identified genera, three genera (*Akkermansia, Eubacterium* and *Lactobacillus*) had positive mediation effects. Genera *Akkermansia* and *Eubacterium* were positively associated with weight gain and STAT increased the weight gain by increasing their relative abundances, while *Lactobacillus* was negatively associated with weight gain and STAT increased the weight gain by decreasing its relative abundance; an unclassified genus from family *Rikenellaceae* had a negative mediation effect: it was positively associated with weight gain, however STAT decreased the weight gain by decreasing its relative abundance. The observation about the effect of *Lactobacillus* on weight gain is consistent with several prior studies (Armougom *et al.*, 2009; Clarke *et al.*, 2012; Turnbaugh *et al.*, 2008).

**Table 2:** The estimated p-values for the microbial mediation effect (at genus rank), the mediation effect estimates and the proportion of the total causal effect on the body weight gain at days 21 and 28 for female and male respectively.

| Day | P-value of |  | Mediation effect |  |
| --- | --- | --- | --- | --- |
|  | OME | CME | Estimate | Proportion* |
| Female |  |  |  |  |
| 21 | 0.154 | 0.438 | 0.102 | 3.0% |
| 28 | 0.048 | 0.040 | 0.634 | 18.8% |
| Male |  |  |  |  |
| 21 | 0.018 | 0.002 | 0.069 | 3.8% |
| 28 | 0.004 | 0.004 | 0.622 | 32.2% |

\* Proportion= ME estimate/TE estimate $\times 100\%$

**Table 3:** Component-wise point and CI estimates of *ME*_*j*_ for the causal genera at day 28 on body weight gain for female mice.

| Genus | Term1* | Term2<sup>§</sup> | $\hat{ME}_j^{\S}$ | 95% $\hat{CI}^{\#}$ | |
| --- | --- | --- | --- | --- | --- |
|  |  |  |  | Lower | Upper |
| *Akkermansia* | 0.118 | 1.466 | 0.173 | 0.126 | 0.220 |
| *Lactobacillus* | -0.301 | -1.289 | 0.388 | 0.324 | 0.452 |
| *Eubacterium* | 0.117 | 1.462 | 0.171 | 0.147 | 0.199 |
| *Rikenellaceae_Other* | 0.065 | -1.508 | -0.098 | -0.119 | -0.077 |

\*Term1 represents $(\alpha_{\hat{M}_j} + \alpha_{\hat{C}_j})$
<sup>§</sup>Term2 represents $\{\hat{E}[\log(M_j)|T = 1] - \hat{E}[\log(M_j)|T = 0]\}$
<sup>§</sup> $\hat{ME}_j = (\alpha_{\hat{M}_j} + \alpha_{\hat{C}_j})\{\hat{E}[\log(M_j)|T = 1] - \hat{E}[\log(M_j)|T = 0]\}$
<sup>#</sup>95% $\hat{CI}$ was calculated by bootstrapping procedure, and the number of bootstrapping is 100.

Similar analyses have been done for male mice. Please see them in the Supplementary Materials, Section S5. In summary, we provide evidence that microbiome plays a significant mediating role in the relationship between antibiotic usage and weight gain for both female and male mice, especially at day 28 in this study. We also observe differences in causal genera between genders, which has been well-demonstrated in Schulfer *et al.* (2019). Among the identified causal genera, the effect directions are mixed and CME test has more significant results than OME, which suggests that CME test is more powerful when both positive and negative component-wise mediation effects are present.

## 4 Discussion

In this paper, we proposed a rigorous causal mediation analytic framework SparseMCMM to investigate the causal mediating role of the high-dimensional and compositional microbiome in the relationship between a treatment and a continuous outcome (see workflow of SparseMCMM in the Supplementary Materials, Section S7). We quantified the causal direct effect of treatment, the overall mediation effect of microbiome community and the component-wise mediation effect for each individual microbe under the counterfactual framework. We developed regularization strategies to handle the high-dimensional mediators and to select the signature causal microbes. We further proposed two tests to examine whether the overall mediation effect of microbiome community on the outcome is significant (test OME) or at least one of component-wise *ME*_*j*_s is significantly non-zero (test CME), respectively. Through extensive simulations, we demonstrated that SparseMCMM provided asymptotically unbiased DE and ME estimates with small MSEs and the proposed tests OME and CME controlled their type I error rate around the significance level even under the constraint of data sparsity. Compared with the competing methods (Delta.T and tau.T), SparseMCMM uniformly achieved higher statistical power under almost all scenarios. Finally, we applied SparseMCMM to analyze a STAT murine microbiome study and to detect unambiguous causal paths among STAT (antibiotic treatment), gut microbiome and body weight (outcome) for both female and male mice respectively.

Recently, Sohn and Li (unpublished at present) proposed a compositional causal mediation model (CMM) to describe the relationships among treatment, microbiome composition, and continuous outcome. Although sharing a similar framework with CMM, our proposed method SparseMCMM has several essential differences. First, we use Dirichlet regression to characterize the relationship between treatment and microbiome composition, while Sohn and Li utilize the algebraic structure of a composition under the simplex space. Secondly, with Dirichlet regression, SparseMCMM can handle the interaction between treatment and microbiome with relation to the outcome in a more flexible manner through the proposed regularization strategy, which addresses concerns regarding the potential bias caused by neglecting the presence of interaction effects (Richiardi *et al.*, 2013; Valeri and VanderWeele, 2013). Moreover, SparseMCMM can automatically drop the interaction terms when the data suggest that they are absent with the proposed penalized least squares criterion; CMM lacks such flexibility. Thirdly, SparseMCMM selects casual taxa with regularization techniques, while CMM identifies the key taxa based on confidence interval estimates.

Considering the completely distinct model assumptions, we did not include CMM in our simulation study. However, we applied CMM (through its R package) to our real data example (only at day 28) to compare its results with ours. For female mice, CMM only identified that genus *Lactobacillus* and an unclassified genus from family *Rikenellaceae* had significant mediation effects on the weight gain. Both of these genera were on the causal list from SparseMCMM. However, no causal overall or component-wise mediation effect was identified by CMM for male mice. Overall, SparseMCMM captured more causal signals than CMM in this real data example, which further demonstrates the sound performance of SparseMCMM.

The phylogenetic tree describes the taxonomical and evolutionary relationships among taxa and provides the possibility to further interpret the casual path among treatment, microbiome and outcome (Knight *et al.*, 2018; Silverman *et al.*, 2017). If the causal taxa are phylogenetically related, utilizing the phylogenetic tree information will increase efficiency of SparseMCMM as in the microbiome association tests (Hu *et al.*, 2018; Koh *et al.*, 2017). Our SparseMCMM R package provides a tree option which takes the prior structure information into account by incorporating a graph Laplacian penalty induced by the phylogenetic tree (Chen *et al.*, 2012; Li and Li, 2008). However, please note that since this option require an additional tuning parameter estimation, it increases the computational complexity. Due to the unknown true nature of the state and the tradeoff of the computational time, we defer the choice of utilization of the phylogenetic tree in SparseMCMM to the user.

In addition, the recent increase in the number of microbial longitudinal studies offers new opportunities to investigate the dynamics of microbial communities. Recent investigations show that the dynamics and stability of the microbial community could be a strong predictor of disease activity (Halfvarson *et al.*, 2017; Novakova *et al.*, 2017), but it is challenging to accommodate the high-dimensional longitudinal mediators in the causal mediation model. One limitation of SparseMCMM is that it can only take a single time point of microbiome data into the proposed framework. As a future research area, we will aim to incorporate microbial dynamic system modelling (Zhang and Davis, 2013) into SparseMCMM to provide a causal path relating treatment, longitudinal microbiome composition, and outcome.

The proposed causal mediation models in this study has been developed into the SparseMCMM R package, and can be installed from https://sites.google.com/site/huilinli09/software and https://github.com/chanw0/SparseMCMM.

