## Supplementary Materials for "Estimating and testing the microbial causal mediation effect with high-dimensional and compositional microbiome data"

#### S1 Identifiability and model assumptions

There are four assumptions must be satisfied in order to identify the causal effect of treatment on outcome using the proposed causal mediation models under the counterfactoral framework (VanderWeele and Vansteelandt, 2009, 2014; VanderWeele, 2016; Huang and Pan, 2016). First, no unmeasured confounders for the relationship between treatment and outcome, i.e., $Y(t, \boldsymbol{m}) \amalg T | \boldsymbol{X}$ for all levels of $t$ and $\boldsymbol{m}$; secondly, no unmeasured confounders for the relationship between mediator and outcome, i.e. $Y(t, \boldsymbol{m}) \amalg \boldsymbol{M} | T, \boldsymbol{X}$ for all levels of $t$ and $\boldsymbol{m}$; thirdly, no unmeasured confounders for the relationship between treatment and mediator, i.e. $\boldsymbol{M}(t) \amalg T | \boldsymbol{X}$ for all levels of $t$; lastly, there is no unmeasured confounders for the relationship between mediator and outcome that can be affected by the treatment, i.e. $Y(t, \boldsymbol{m}) \amalg \boldsymbol{M}(t^*) | \boldsymbol{X}$ for all levels of $t$, $t^*$ and $\boldsymbol{m}$.

#### S2 The derivation of the causal DE and ME estimators

Under the four sufficient identifiability assumptions discussed in Section S1 and the models (1) and (2) in the manuscript, the causal direct effect of treatment on the outcome is defined as

$$
\begin{aligned}
DE &= E[Y_{T=1,\boldsymbol{M}(T=0)} - Y_{T=0,\boldsymbol{M}(T=0)}|\boldsymbol{X}] \\
&= \int [E(Y|T=1,\boldsymbol{M},\boldsymbol{X}) - E(Y|T=0,\boldsymbol{M},\boldsymbol{X})]dF(\boldsymbol{M}|T=0,\boldsymbol{X}) \\
&= \int \Big\{\alpha_0 + \alpha_T + \boldsymbol{\alpha}_X^T\boldsymbol{X} + \boldsymbol{\alpha}_M^T[\log(\boldsymbol{M})] + \boldsymbol{\alpha}_C^T\log(\boldsymbol{M}) - \alpha_0 - \boldsymbol{\alpha}_X^T\boldsymbol{X} - \\
&\boldsymbol{\alpha}_M^T[\log(\boldsymbol{M})]\Big\}dF(\boldsymbol{M}|T=0,\boldsymbol{X}) \\
&= \int \left\{\alpha_T + \boldsymbol{\alpha}_C^T[\log(\boldsymbol{M})]\right\} dF(\boldsymbol{M}|T=0,\boldsymbol{X}) \\
&= \alpha_T + \boldsymbol{\alpha}_C^T E[\log(\boldsymbol{M})|T=0,\boldsymbol{X}].
\end{aligned}
$$

The mediation effect of microbiome composition on the outcome is defined as

$$
\begin{aligned}
ME &= E[Y_{T=1,\boldsymbol{M}(T=1)} - Y_{T=1,\boldsymbol{M}(T=0)}|\boldsymbol{X}] \\
&= \int E(Y|T=1,\boldsymbol{M},\boldsymbol{X})dF(\boldsymbol{M}|T=1,\boldsymbol{X}) - \int E(Y|T=1,\boldsymbol{M},\boldsymbol{X})dF(\boldsymbol{M}|T=0,\boldsymbol{X}) \\
&= \int \left\{\alpha_0 + \alpha_T + \boldsymbol{\alpha}_X^T\boldsymbol{X} + \boldsymbol{\alpha}_M^T[\log(\boldsymbol{M})] + \boldsymbol{\alpha}_C^T\log(\boldsymbol{M})\right\} dF(\boldsymbol{M}|T=1,\boldsymbol{X}) - \\
&\int \left\{\alpha_0 + \alpha_T + \boldsymbol{\alpha}_X^T\boldsymbol{X} + \boldsymbol{\alpha}_M^T[\log(\boldsymbol{M})] + \boldsymbol{\alpha}_C^T\log(\boldsymbol{M})\right\} dF(\boldsymbol{M}|T=0,\boldsymbol{X}) \\
&= (\boldsymbol{\alpha}_M^T + \boldsymbol{\alpha}_C^T)\{E[\log(\boldsymbol{M})|T=1,\boldsymbol{X}] - E[\log(\boldsymbol{M})|T=0,\boldsymbol{X}]\}.
\end{aligned}
$$

Therefore, the total treatment effect on the outcome is

$$
\begin{aligned}
TE &= DE + ME \\
&= E[Y_{T=1,\boldsymbol{M}(T=1)} - Y_{T=0,\boldsymbol{M}(T=0)}|\boldsymbol{X}] \\
&= \alpha_T + \boldsymbol{\alpha}_C^T E[\log(\boldsymbol{M})|T=0,\boldsymbol{X}] + (\boldsymbol{\alpha}_M^T + \boldsymbol{\alpha}_C^T)\{E[\log(\boldsymbol{M})|T=1,\boldsymbol{X}] - E[\log(\boldsymbol{M})|T=0,\boldsymbol{X}]\} \\
&= \alpha_T + (\boldsymbol{\alpha}_M^T + \boldsymbol{\alpha}_C^T)\{E[\log(\boldsymbol{M})|T=1,\boldsymbol{X}] - \boldsymbol{\alpha}_M^T E[\log(\boldsymbol{M})|T=0,\boldsymbol{X}].
\end{aligned}
$$

Here $E[\log(\boldsymbol{M})|T=0,\boldsymbol{X}]$ is the conditional expectation of $\log(\boldsymbol{M})$ given $T=0$ and covariates $\boldsymbol{X}$, where $\boldsymbol{M}$ follows the Dirichlet distribution with parameters $\boldsymbol{\gamma}=(\gamma_1,\ldots,\gamma_p)^T$. Due to the aggregation property of Dirichlet distribution, $(M_j, 1-M_j)$ follows the two dimensional Dirichlet distribution as Dirichlet$(\gamma_j, \sum_{j'=1}^{p}\gamma_{j'}-\gamma_j)$ given $T=0$ and covariates $\boldsymbol{X}$. Then $E[\log(M_j)|T=0,\boldsymbol{X}]$ can be calculated as the following (Honkela, 2001):

$$
\begin{aligned}
E[\log(M_j)|T=0,\boldsymbol{X}] &= \int_0^1 \frac{\Gamma\left(\sum_{j'=1}^{p}\gamma_{j'}\right)}{\Gamma(\gamma_j)\Gamma\left(\sum_{j'=1}^{p}\gamma_{j'}-\gamma_j\right)} M^{\gamma_j-1}(1-M)^{\sum_{j'=1}^{p}\gamma_{j'}-\gamma_j-1}\log M dM \\
&= \psi\left[\gamma_j(T=0,\boldsymbol{X})\right]-\psi\left[\sum_{j'=1}^{p}\gamma_{j'}(T=0,\boldsymbol{X})\right],
\end{aligned}
$$

where $\gamma_j(T=0,\boldsymbol{X})=\exp\left(\beta_{0j}+\boldsymbol{\beta}_{\boldsymbol{X}j}^T\boldsymbol{X}\right)$ and $\psi(\cdot)=\frac{d}{dx}\ln(\Gamma(x))$ is the digamma function. Similarly, $E[\log(M_j)|T=1,\boldsymbol{X}]=\psi\left[\gamma_j(T=1,\boldsymbol{X})\right]-\psi\left[\sum_{j'=1}^{p}\gamma_{j'}(T=1,\boldsymbol{X})\right]$. Thus we have

$$
\begin{aligned}
E[\log(\boldsymbol{M})|T=0,\boldsymbol{X}] = \Bigg(&\psi\left[\gamma_1(T=0,\boldsymbol{X})\right]-\psi\left[\sum_{j=1}^{p}\gamma_j(T=0,\boldsymbol{X})\right],\ldots, \\
&\psi\left[\gamma_p(T=0,\boldsymbol{X})\right]-\psi\left[\sum_{j=1}^{p}\gamma_j(T=0,\boldsymbol{X})\right]\Bigg)^T, \text{ and} \\
E[\log(\boldsymbol{M})|T=1,\boldsymbol{X}] = \Bigg(&\psi\left[\gamma_1(T=1,\boldsymbol{X})\right]-\psi\left[\sum_{j=1}^{p}\gamma_j(T=1,\boldsymbol{X})\right],\ldots, \\
&\psi\left[\gamma_p(T=1,\boldsymbol{X})\right]-\psi\left[\sum_{j=1}^{p}\gamma_j(T=1,\boldsymbol{X})\right]\Bigg)^T,
\end{aligned}
$$

where $\gamma_j(T=0,\boldsymbol{X})=\exp\left(\beta_{0j}+\boldsymbol{\beta}_{Xj}^T\boldsymbol{X}\right)$, $\gamma_j(T=1,\boldsymbol{X})=\exp\left(\beta_{0j}+\beta_{Tj}+\boldsymbol{\beta}_{Xj}^T\boldsymbol{X}\right)$, $j=1,\ldots,p$, and $\psi(\cdot)=\frac{d}{dx}\ln(\Gamma(x))$ is the digamma function.

### S3 Competing methods

#### S3.1 Delta.T and tau.T tests

Huang and Pan (2016) developed a causal mediation model particularly focusing on testing the overall mediation effect for the high-dimensional continuous mediators with application in gene expression studies. Specifically, two multivariate linear regressions are proposed to describe the relationships among outcome, treatment and mediators, as well as, the relationship of treatment and mediators respectively. In their first multivariate linear regression, the outcome $Y_i$ is determined by $q$ covariates $\boldsymbol{X}_i$, one treatment $T_i$, $p$ mediators $\mathbf{G}_i$ and interactions between treatment $T_i$ and mediators $\mathbf{G}_i$.

$$Y_i = \alpha_0 + \boldsymbol{\alpha}_X^T \boldsymbol{X}_i + \alpha_T T_i + \boldsymbol{\alpha}_G^T \mathbf{G}_i + \boldsymbol{\alpha}_C^T \boldsymbol{G}_i T_i + \epsilon_{Yi}, \tag{S1}$$

where $\alpha_0$ is the intercept, $\alpha_T$ is the coefficient of treatment, $\boldsymbol{\alpha}_X = (\alpha_{X1}, \ldots, \alpha_{Xq})^T$, $\boldsymbol{\alpha}_G = (\alpha_{G1}, \ldots, \alpha_{Gp})^T$, and $\boldsymbol{\alpha}_C = (\alpha_{C1}, \ldots, \alpha_{Cp})^T$ are the vectors of coefficients of covariates, mediators, and interactions between treatment and mediators, respectively, and $\epsilon_{Yi} \sim N(0, \sigma^2)$ is the error term.

Further, mediators are determined by $q$ covariates and treatment in their second regression.

$$\mathbf{G}_i = \boldsymbol{\beta}_0 + \mathbf{B}\boldsymbol{X}_i + T_i \boldsymbol{\beta}_T + \boldsymbol{\epsilon}_{Gi}, \tag{S2}$$

where $\boldsymbol{\beta}_0^T = (\beta_{01}, \ldots, \beta_{0p})$, $\mathbf{B}^T = (\boldsymbol{\beta}_{X1}, \ldots, \boldsymbol{\beta}_{Xp})$, $\boldsymbol{\beta}_T^T = (\beta_{T1}, \ldots, \beta_{Tp})$ and $\boldsymbol{\epsilon}_{Gi} \sim N_p(\mathbf{0}, \Sigma^2)$.

With four identifiable assumptions, they define the mediation effect with the parameters in models (S1) and (S2) as:

$$E[Y_{T=1,\boldsymbol{G}(T=1)} - Y_{T=1,\boldsymbol{G}(T=0)} | \boldsymbol{X}] = \boldsymbol{\beta}_T^T(\boldsymbol{\alpha}_G + \boldsymbol{\alpha}_C). \tag{S3}$$

Obviously, Equation (S3) can only evaluate mediation effect for the dataset with sample size $n$ larger than the number of mediators $p$. To accommodate the setting with a larger number of mediators and a small sample size, Huang and Pan (2016) constructed a transformed model by utilizing the spectral decomposition on the covariance matrix of correlated mediators to reduce the dimensionality of the mediators, as mentioned in the Introduction. Then their mediation effect is expressed as

$$E[Y_{T=1,\boldsymbol{P}(T=1)} - Y_{T=1,\boldsymbol{P}(T=0)}|\boldsymbol{X}] = \boldsymbol{\beta}_T^{*T}(\boldsymbol{\alpha}_G^* + \boldsymbol{\alpha}_C^*), \tag{S4}$$

where, $\boldsymbol{\beta}_0^* = \mathbf{u}\boldsymbol{\beta}_0$, $\boldsymbol{\beta}_T^* = \mathbf{u}\boldsymbol{\beta}_T$, $\boldsymbol{\alpha}_G^* = \mathbf{u}\boldsymbol{\alpha}_G$, $\boldsymbol{\alpha}_C^* = \mathbf{u}\boldsymbol{\alpha}_C$. $\mathbf{u}$ is an orthogonal matrix such that $\mathbf{u}\Sigma\mathbf{u}^T = \text{diag}(\sigma_1^2, \ldots, \sigma_p^2)$.

They proposed three tests to test the overall mediation effect based on equation (S4), as the following respectively

$$\begin{aligned}
&H_{10} : \text{Delta.T} \equiv \boldsymbol{\beta}_T^{*T}(\boldsymbol{\alpha}_G^* + \boldsymbol{\alpha}_C^*) = 0\\
&H_{20} : \delta_j \equiv \boldsymbol{\beta}_{Tj}^{*T}(\boldsymbol{\alpha}_{Gj}^* + \boldsymbol{\alpha}_{Cj}^*) = 0, \quad \forall j \in 1, \ldots, p\\
&H_{30} : \text{tau.T} \equiv \left[\boldsymbol{\beta}_{Tj}^{*T}(\boldsymbol{\alpha}_{Gj}^* + \boldsymbol{\alpha}_{Cj}^*)\right]^2 = 0.
\end{aligned}$$

Specifically, Delta.T tests whether the overall mediation effect of all mediators exists or not. The last two are equivalent and test whether at least one mediator has a significant mediation effect. Similar tests for the original mediators $\mathbf{G}$ based on equation (S3) were also proposed. Since the setting with a large $p$ and a small $n$ is our interest, Delta.T and tau.T, which have been demonstrated to have superior performance in Huang and Pan (2016), are used in our comparisons. Moreover, the normality-based and bootstrap-based methods are used to approximate the distributions of Delta.T and tau.T with Monte-Carlo procedure, respectively, to evaluate their testing performance.

#### S3.2 HIMA test

Zhang *et al.* (2016) extended the multiple mediator models (Preacher and Hayes, 2008) to the high-dimensional setting and proposed a test (denoted as HIMA) to test mediation effects at the individual-level in high-dimensional epigenetic studies. The relationships among outcome, treatment and mediators in their method are described as below

$$Y_i = \alpha_0 + \alpha_T T_i + \boldsymbol{\alpha}_G^T \mathbf{G}_i + \epsilon_{Y_i}, \tag{S5}$$

$$G_k = \beta_{0k} + \beta_k T_i + \epsilon_{G_k}, k = 1, \ldots, p, \tag{S6}$$

where $\alpha_T$ and $\boldsymbol{\alpha}_G = (\alpha_{G1}, \ldots, \alpha_{Gp})^T$ are the coefficients of treatment and mediators on the outcome respectively. $\beta_k$ is the coefficient of treatment on the $k$the mediator, and the mediation effect for the $k^{\text{th}}$ mediator is $\alpha_{Gk}\beta_k$, $k = 1, \ldots, p$.

Since traditional regression analysis fails to give the estimates $\hat{\alpha}_{Gk}$ $k = 1, \ldots, p$ based on the regression (S5) in the high-dimensional setting (the number of mediators $p$ is much larger than the sample size), they reduce the dimension of mediators by first using the sure independent screening (Fan and Lv, 2008) to identify a subset of mediators with a moderate number (less than sample size) and then employing minimax concave penalty techniques (Zhang *et al.*, 2010) to determine selected mediators. Then they obtain p values $p_k^\alpha$ and $p_k^\beta$ for the $k$th selected mediators by testing $H_0 : \alpha_{Gk} = 0$ and $H_0 : \beta_k = 0$ with Wald test based on regressions (S5)-(S6) respectively. Finally they define the p values for testing mediation effect of the $k$th selected mediator as $\max(p_k^\alpha, p_k^\beta)$. Here they use Bonferroni's method to adjust for multiple comparisons.

Obviously, none of the tests in Section S3.1 and S3.2 is designed to deal with compositional microbiome data. In the simulation studies and real data analysis, we log transformed the relative abundances to make them more normal before using those three tests to test microbial mediation effects. In addition, since Delta.T and tau.T do not satisfy multivariate normality assumption because of the constraint $\sum_{ij}^{p} M_{ij} = 1$, $i = 1, \ldots, n$. So we adopt their bootstrap-based method to calculate their p-values. For HIMA test, we calculated its type I error rate and power in simulation studies following Dezeure *et al.* (2015) and Zhang *et al.* (2016).

#### S4 Bootstrapping procedure for CI estimation

We use the following bootstrapping procedure to calculate the approximated 95% confidence intervals (CIs) of the component-wise $ME_j$ estimates for all causal genera.

**Step 1**: Randomly sample $n$ subjects with replacement from the original data $(T_i, \boldsymbol{X}_i, \boldsymbol{M}_i, Y_i)$, $i = 1, \ldots, n$, denoted as $(T_{(i)}, \boldsymbol{X}_{(i)}, \boldsymbol{M}_{(i)}, Y_{(i)})$, $i = 1, \ldots, n$.

**Step 2**: Given the sampled data $(T_{(i)}, \boldsymbol{X}_{(i)}, \boldsymbol{M}_{(i)}, Y_{(i)})$, $i = 1, \ldots, n$, we estimate the coefficients of all parameters in equations (3)-(4), and then calculate the component-wise $ME_j$ estimates.

**Step 3**: Repeat Steps 1-2 $R$ times to get $R$ sets of the component-wise $ME_j$ estimates $\left\{ \left( \hat{ME}_j^{(r)} \right), j = 1, \ldots, p \right\}_{r=1}^{R}$.

**Step 4**: Based on $\left\{ \left( \hat{ME}_j^{(r)} \right), j = 1, \ldots, p \right\}_{r=1}^{R}$, calculate the corresponding standard deviation (SD) for all genera, $\hat{SD}_j$, $j = 1, \ldots, p$.

**Step 5**: The approximated 95% CI estimate of $ME_j$ is $\hat{CI}_j = \hat{ME}_j \pm \frac{1.96 \times \hat{SD}_j}{\sqrt{R}}$, where $\hat{ME}_j$ is calculated based on the original data, $j = 1, \ldots, p$.

#### S5 Analyses for the male mice in the real data analysis

Similar analyses have been done for the male mice. In **Table 3**, both OME and CME tests are significant at both time points. The estimated overall MEs are 0.069 and 0.622, which represent 3.6% and 32.2% of the total causal treatment effect on the weight gain at days 21 and 28, respectively. Note that the overall ME estimate at day 21 is weak and the estimated non-zero component-wise $ME_j$s are also very small (**Table S10**), so we focus on day 28 in the following discussion. Compared to the females, the microbiome of male mice exhibits a much stronger mediation effect with much smaller p values at day 28. Further, SparseMCMM detected six causal genera at day 28 reported in **Table S10** with their ME and 95% CI estimates. However since the last two genera have very small ME estimates and their CI estimates include zero, they are not reported as the significant causal taxa.

Among the four identified genera: STAT has a positive effect on the weight gain through *Oscillospira*, an unclassified genus from family *Rikenellaceae* and an unclassified genus from order *Clostridiales*, while STAT has a negative effect on weight gain through *Bifidobacterium* (**Table S11**). Several other murine studies have shown that higher abundance of *Oscillospira* in the gut microbiota was associated with lower body mass index and has been regarded as a probiotic bacteria in the digestive system (Stenman *et al.*, 2016; Goodrich *et al.*, 2014; Konikoff and Gophna, 2016), which agree with the findings here.

#### S6 Availability of data and materials

The murine microbiome data (Schulfer *et al.*, 2019) we used in the real data analysis are available at QIITA database (`https://qiita.ucsd.edu`) and the European Bioinformatics Institute (EBI) database (`https://www.ebi.ac.uk`), under the project ID PRJEB18627 (`http://www.ebi.ac.uk/ena/data/view/PRJEB18627`).

The R package SparseMCMM is publicly available at `https://sites.google.com/site/huilinli09/software` and `https://github.com/chanw0/SparseMCMM`.

#### S7 Workflow of SparseMCMM

In this section, we give a workflow of SparseMCMM to clarify its methods, procedures and delivery outputs as illustrated in **Fig. S1**. SparseMCMM is designed for the high dimensional and compositional microbiome data in a treatment-microbiome-outcome causal study. SparseMCMM utilizes the linear log-contrast regression and Dirichlet regression to quantify the causal direct effect of the treatment and the causal mediation effect of the microbiome on the outcome under the counterfactual framework while addressing the compositional structure of microbiome data. Further it implements regularization techniques to handle the high-dimensional microbial mediators and identify the signature causal microbes.

The workflow of SparseMCMM consists of three steps:

**Step 1**: Report the estimates of DE, ME and TE respectively under the sufficient causal assumptions mentioned in the Supplementary Materials, Section S1.

**Step 2**: Report the overall mediation test results: OME test can determine whether the overall mediation effect of microbiome is significant, and CME test can determine whether at least one individual microbe has a significant mediation effect on the outcome.

**Step 3**: Report the point and 95% confidence interval estimates of $ME_j$ for each signature causal microbe identified by the regularization technique only if CME test is significant at step 2.

Consequentially, SparseMCMM provides a clear and sensible causal path analysis among treatment, compositional microbiome and outcome.

#### S8 Supplementary tables

Table S1: The log-transformed baseline relative abundance $\boldsymbol{\beta}_0$ used in the simulation studies

| $p$ | $\boldsymbol{\beta}_0$ |
|---|---|
| 10 | $(0.6, -0.3, 0.8, -1.4, -1.2, -1.4, -1.3, -1.0, -0.2, 0.6)^T$ |
| 25 | $(1.6, -1.4, -0.8, -2.5, -2.2, -1.3, -1.1, -0.3, -1.9, -1.7, -4.6, -0.2, -1.6,$ $-3.0, -2.3, -3.9, -1.4, -1.4, -2.2, -2.7, -3.8, -2.8, -2.8, -0.5, -2.8)^T$ |
| 50 | $(-2.3, -0.2, -2.5, -1.3, -1.9, -1.9, -0.4, 0.3, -4.4, -3.7, 0.1, -1.3, -3.7, -1.9,$ $-4.4, -3.2, -3.7, -2.8, -3.7, -1.8, -3.1, -0.4, -0.9, -4.4, -1.8, -4.4, -2.5, -3.7,$ $-4.4, -3.7, -2.8, -3.7, -3.7, -4.4, -0.5, -1.5, -3.7, -4.4, -0.7, -3.3, -2.1, -4.4,$ $-0.7, -3.7, -4.4, -4.4, -1.4, -1.8, -1.8, -2.3)^T$ |

Table S2: Parameter settings used for estimating DE and ME in the simulation studies

| | | Indices of the | Regression coefficients for the causal taxa | | |
|---|---|---|---|---|---|
| $p$ | $p_r$ | causal taxa | $\boldsymbol{\alpha}_M^*$ | $\boldsymbol{\alpha}_C^*$ | $\boldsymbol{\beta}_T^*$ |
| 10 | 2 | 1,3 | $(-0.45, 0.45)^T$ | $(-0.82, 0.82)^T$ | $(0.26, 0.65)^T$ |
| 25 | 3 | 2,8,13 | $(0.83, -0.055, -0.78)^T$ | $(0.56, -0.08, -0.48)^T$ | $(1.95, 1.95, 1.35)^T$ |
| 50 | 5 | 2,7,11,22,35 | $(1.1, -1.47, 0.81, -0.16, -0.29)^T$ | $(-0.39, -0.93, 0.50, 0.15, 0.67)^T$ | $(-1.38, 0.01, 2, 0.18, 0.68)^T$ |

Table S3: True TEs and MEs based on the parameter settings in Table S2

| $p$ | $p_r$ | Indices of the causal taxa | TE | ME | Component-wise $ME_j$s for the causal taxa |
|---|---|---|---|---|---|
| 10 | 2 | 1,3 | 1.766 | 0.555 | -0.081, 0.636 |
| 25 | 3 | 2,8,13 | 1.808 | 0.605 | 5.570, -0.290, -4.675 |
| 50 | 5 | 2,7,11,22,35 | 1.796 | 0.642 | -3.396, 1.318, 2.469, 0.002, 0.249 |

Table S4. Computational time in the estimating part (in seconds)

| Number of mediators | Sample size $n$ | | | |
|---|---|---|---|---|
| $p$ | 50 | 100 | 300 | 500 |
| 10 | 1.75 | 2.27 | 3.07 | 3.21 |
| 25 | 15.59 | 14.24 | 26.64 | 36.22 |
| 50 | 141.95 | 169.00 | 199.50 | 250.68 |
| 100 | 320.40 | 365.4 | 514.20 | 581.40 |

Table S5: Parameter settings used for hypothesis testing in the simulation studies

| | | | Quantification | | Regression coefficients for the causal taxa | | |
|---|---|---|---|---|---|---|---|
| $p$ | Scenario | Qualification | ME | CME | $\boldsymbol{\alpha}_M^*$ | $\boldsymbol{\alpha}_C^*$ | $\boldsymbol{\beta}_T^*$ |
| 10 | 1 | null | 0 | 0 | $(0,0)^T$ | $(0,0)^T$ | $(0,0)^T$ |
| | | small | 0.29 | 0.04 | $(0.35,-0.35)^T$ | $(0.1,-0.1)^T$ | $(0.3,-0.2)^T$ |
| | | medium | 0.51 | 0.13 | $(0.7,-0.7)^T$ | $(0.1,-0.1)^T$ | $(0.3,-0.2)^T$ |
| | | large | 0.70 | 0.25 | $(1.0,-1.0)^T$ | $(0.1,-0.1)^T$ | $(0.3,-0.2)^T$ |
| 10 | 2 | null | 0 | 0 | $(0,0)^T$ | $(0,0)^T$ | $(0,0)^T$ |
| | | small | 0.26 | 0.13 | $(0.7,-0.7)^T$ | $(0.3,-0.3)^T$ | $(0.4,0.2)^T$ |
| | | medium | 0.39 | 0.30 | $(0.9,-0.9)^T$ | $(0.6,-0.6)^T$ | $(0.4,0.2)^T$ |
| | | large | 0.52 | 0.52 | $(1.2,-1.2)^T$ | $(0.8,-0.8)^T$ | $(0.4,0.2)^T$ |
| 25 | 1 | null | 0 | 0 | $(0,0,0)^T$ | $(0,0,0)^T$ | $(0,0,0)^T$ |
| | | small | 0.43 | 0.07 | $(0.2,-0.3,0.1)^T$ | $(0.1,-0.2,0.1)^T$ | $(0.1,-0.2,0.1)^T$ |
| | | medium | 0.60 | 0.13 | $(0.3,-0.5,0.2)^T$ | $(0.1,-0.2,0.1)^T$ | $(0.1,-0.2,0.1)^T$ |
| | | large | 1.18 | 0.59 | $(0.6,-0.9,0.3)^T$ | $(0.1,-0.2,0.1)^T$ | $(0.1,-0.3,0.1)^T$ |
| 25 | 2 | null | 0 | 0 | $(0,0,0)^T$ | $(0,0,0)^T$ | $(0,0,0)^T$ |
| | | small | 0.45 | 0.37 | $(0.2,-0.4,0.2)^T$ | $(0.3,-0.5,0.2)^T$ | $(0.2,0.2,0.2)^T$ |
| | | medium | 0.63 | 2.73 | $(0.7,-1,0.3)^T$ | $(0.4,-0.6,0.2)^T$ | $(0.3,0.4,0.2)^T$ |
| | | large | 1.13 | 6.15 | $(0.9,-1.3,0.4)^T$ | $(0.4,-0.6,0.2)^T$ | $(0.4,0.5,0.3)^T$ |
| 50 | 1 | null | 0 | 0 | $(0,0,0,0,0)^T$ | $(0,0,0,0,0)^T$ | $(0,0,0,0,0)^T$ |
| | | small | 0.84 | 0.17 | $(0.3,0.6,-0.3,0.2,-0.8)^T$ | $(0.2,0.1,-0.3,0.2,-0.2)^T$ | $(0.2,0.2,-0.1,0.1,-0.1)^T$ |
| | | medium | 1.40 | 0.50 | $(0.4,0.9,-0.6,0.2,-0.9)^T$ | $(0.2,0.1,-0.3,0.2,-0.2)^T$ | $(0.2,0.2,-0.1,0.2,-0.2)^T$ |
| | | large | 1.78 | 1.25 | $(0.4,0.9,-0.3,0.2,-1.2)^T$ | $(0.2,0.1,-0.3,0.2,-0.2)^T$ | $(0.2,0.2,-0.1,0.1,-0.3)^T$ |
| 50 | 2 | null | 0 | 0 | $(0,0,0,0,0)^T$ | $(0,0,0,0,0)^T$ | $(0,0,0,0,0)^T$ |
| | | small | 0.67 | 0.90 | $(1,0.8,-0.8,0.7,-1.7)^T$ | $(0.2,0.1,-0.3,0.2,-0.2)^T$ | $(0.2,0.2,-0.1,0.3,0.2)^T$ |
| | | medium | 1.00 | 2.11 | $(1,1.5,-1.5,0.7,-1.7)^T$ | $(0.2,0.1,-0.3,0.2,-0.2)^T$ | $(0.2,0.2,-0.2,0.3,0.3)^T$ |
| | | large | 1.43 | 4.53 | $(1.3,1.5,-2,1,-1.8)^T$ | $(0.2,0.1,-0.3,0.2,-0.2)^T$ | $(0.2,0.2,-0.3,0.3,0.4)^T$ |

Table S6. Computational time in the testing part. Sample size=100 and permutation=500.

| $p$ | 10 | 25 | 50 | 100 |
|---|---|---|---|---|
| Minutes | 1.02 | 7.83 | 36.89 | 61.80 |

Table S7: Empirical Type I error rates of the proposed tests: OME and CME; and the competing tests: Delta.T, tau.T and HIMA for various dimensions of the compositional mediators when the sample size $n = 100$ (significance level=5%).

| $p$ | OME | CME | Delta.T | tau.T | HIMA |
|---|---|---|---|---|---|
| 10 | 1.2% | 1.2% | 0.6% | 0.4% | 0.8% |
| 25 | 0.9% | 1.5% | 0.5% | 0.3% | 1.9% |
| 50 | 0.6% | 1.9% | 0.2% | 0.4% | 2.3% |

Table S8: The estimated p-values of Delta.T and tau.T for the microbial mediation effect (at genus rank) on the body weight gain at days 21 and 28 for female and male respectively.

| | | P-value of | |
|---|---|---|---|
| Sex | Day | Delta.T | tau.T |
| Female | 21 | 0.120 | 0.602 |
| | 28 | 0.072 | 0.261 |
| Male | 21 | 0.846 | 0.830 |
| | 28 | 0.798 | 0.623 |

Table S9: The estimated p-values of the selected genera based on HIMA for the microbial mediation effect on the body weight gain at days 21 and 28 for female and male respectively.

| Female | | | |
|---|---|---|---|
| Day 21 | | Day 28 | |
| Genus | P-value | Genus | P-value |
| *Turicibacter* | 0.802 | *Clostridiaceae_Other* | 0.145 |
| *Coriobacteriaceae_Other* | 1 | *Lactobacillus* | 0.589 |
| *Dehalobacterium* | 1 | *Coriobacteriaceae_Other* | 1 |
| *RF39_Other* | 1 | *Streptophyta_Other* | 1 |
| *Coprococcus* | 1 | | |
| *Ruminococcus* | 1 | | |
| *Allobaculum* | 1 | | |
| *Oscillospira* | 1 | | |
| *Adlercreutzia* | 1 | | |
| *Clostridiales_Other* | 1 | | |
| Male | | | |
| Day 21 | | Day 28 | |
| Genus | P-value | Genus | P-value |
| *Candidatus Arthromitus* | 0.002 | *Ruminococcus* | 0.014 |
| *Allobaculum* | 0.089 | *Clostridium* | 0.568 |
| *Enterobacteriaceae_Other* | 0.504 | *Streptophyta_Other* | 1 |
| *Clostridium* | 0.626 | *S24-7_Other* | 1 |
| *Odoribacter* | 1 | *Rikenellaceae_Other* | 1 |
| | | *Bifidobacterium* | 1 |
| | | *Ruminococcaceae_Other* | 1 |
| | | *Enterobacteriaceae_Other* | 1 |

Table S10: Component-wise point and CI estimates of $ME_j$ for the causal genera at day 21 on body weight gain for male mice.

| Genus | Term1* | Term2$ | $\hat{ME}_j^{\S}$ | 95% $\hat{CI}^{\#}$ Lower | Upper |
|---|---|---|---|---|---|
| *Candidatus Arthromitus* | -0.029 | -1.414 | 0.041 | 0.006 | 0.076 |
| *Enterobacteriaceae_Other* | 0.073 | 0.219 | 0.016 | 0.001 | 0.030 |
| *S24-7_Other* | -0.040 | -0.325 | 0.013 | 0.009 | 0.018 |
| *Akkermansia* | -0.004 | 0.0225 | -9.00E-04 | -0.005 | 0.003 |

*Term1 represents $(\hat{\alpha_{Mj}} + \hat{\alpha_{Cj}})$

$Term2 represents $\{\hat{E}[\log(M_j)|T = 1] - \hat{E}[\log(M_j)|T = 0]\}$

§$\hat{ME}_j = (\hat{\alpha_{Mj}} + \hat{\alpha_{Cj}})\{\hat{E}[\log(M_j)|T = 1] - \hat{E}[\log(M_j)|T = 0]\}$

#95% $\hat{CI}$ was calculated by bootstrapping procedure, and the number of bootstrapping is 100.

Table S11: Component-wise point and CI estimates of $ME_j$ for the causal genera at day 28 on body weight gain for male mice.

| | | | | 95% $\hat{CI}^{\#}$ | |
|---|---|---|---|---|---|
| Genus | Term1* | Term2$ | $\hat{ME}_j^{\S}$ | Lower | Upper |
| *Oscillospira* | -0.124 | -1.492 | 0.185 | 0.156 | 0.214 |
| *Rikenellaceae_Other* | 0.074 | 0.500 | 0.037 | 0.029 | 0.045 |
| *Clostridiales_Other_Other* | -0.235 | -1.736 | 0.408 | 0.366 | 0.450 |
| *Bifidobacterium* | 0.267 | -0.067 | -0.018 | -0.035 | -0.001 |
| *Eubacterium* | 0.007 | 0.571 | 0.004 | 0.000 | 0.008 |
| *Enterobacteriaceae_Other* | 0.010 | 0.060 | 0.006 | -0.005 | 0.017 |

*Term1 represents $(\hat{\alpha_{Mj}} + \hat{\alpha_{Cj}})$

$Term2 represents $\{\hat{E}[\log(M_j)|T = 1] - \hat{E}[\log(M_j)|T = 0]\}$

§$\hat{ME}_j = (\hat{\alpha_{Mj}} + \hat{\alpha_{Cj}})\{\hat{E}[\log(M_j)|T = 1] - \hat{E}[\log(M_j)|T = 0]\}$

#95% $\hat{CI}$ was calculated by bootstrapping procedure, and the number of bootstrapping is 100.

#### S9 Supplementary figures

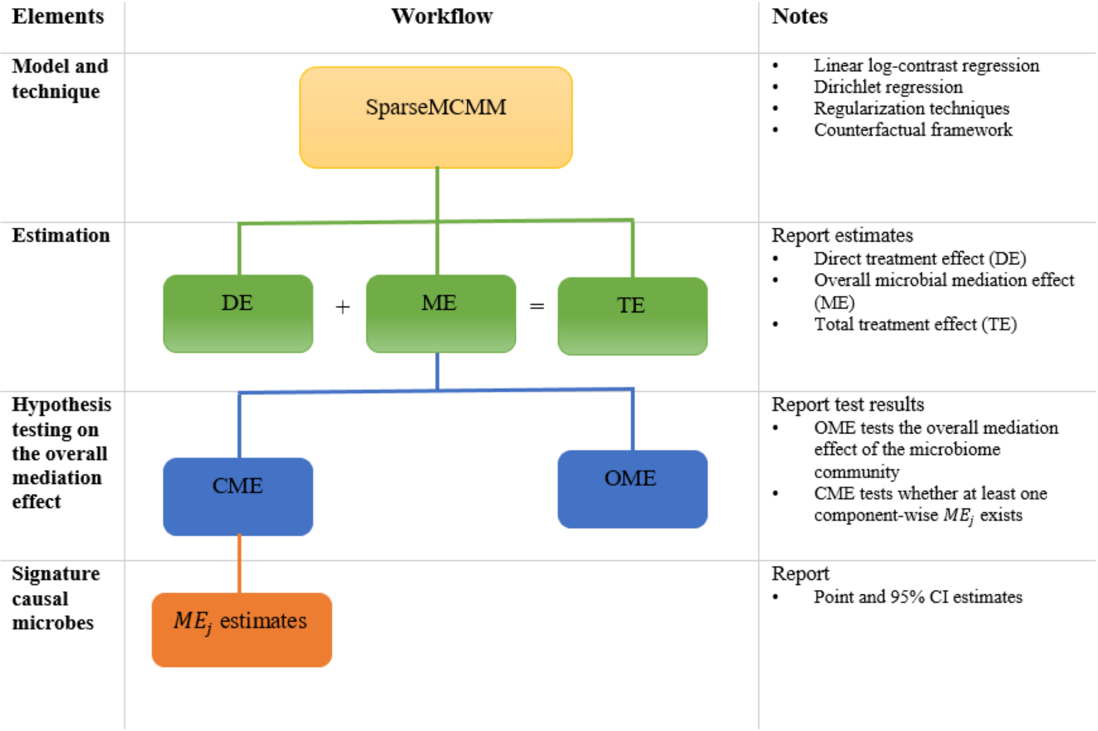


Fig. S1. Workflow of SparseMCMM.

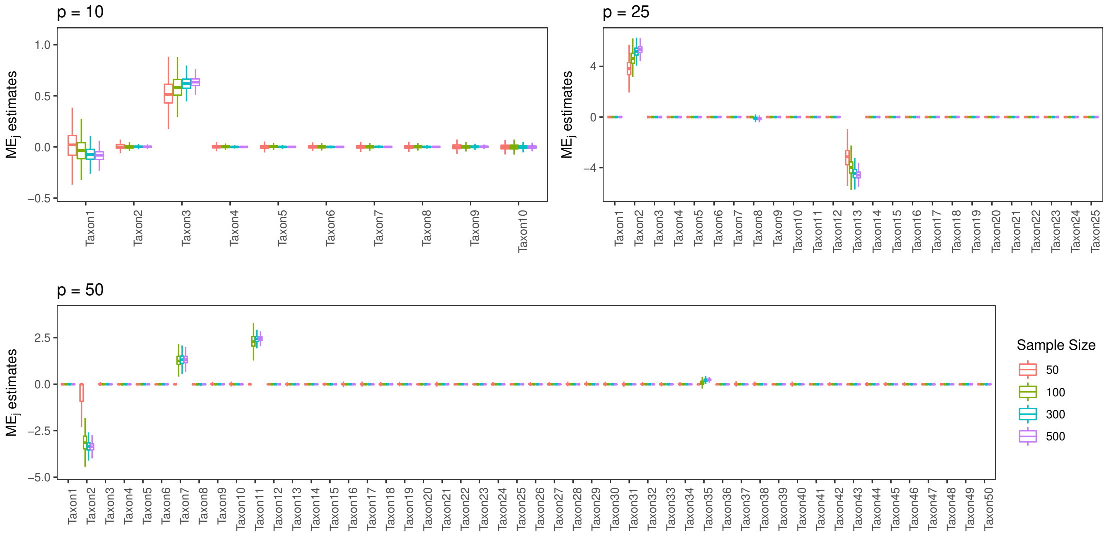


Fig. S2. The component-wise $ME_j$ estimates with sample size $n = 50, 100, 300$ and $500$ respectively. The number of compositional mediators is $p = 10, 25$ and $50$ respectively.

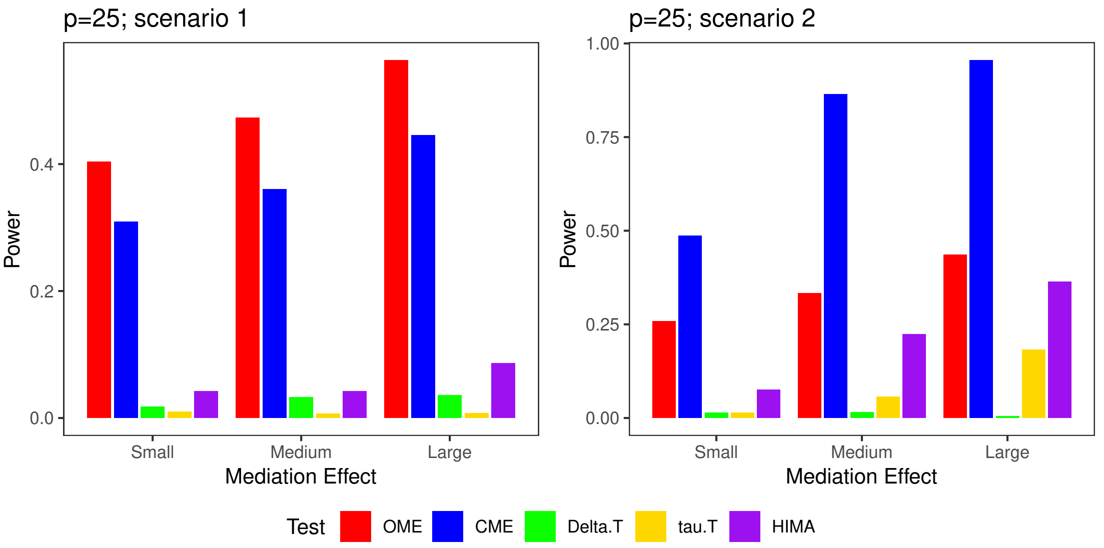


Fig. S3. Empirical power for testing mediation effect with $p = 25$ in scenarios 1-2 (significance level=5%). Note that the magnitudes of mediation effect are not comparable across different $p$s and different scenarios. The detailed setting is given in Table S5.

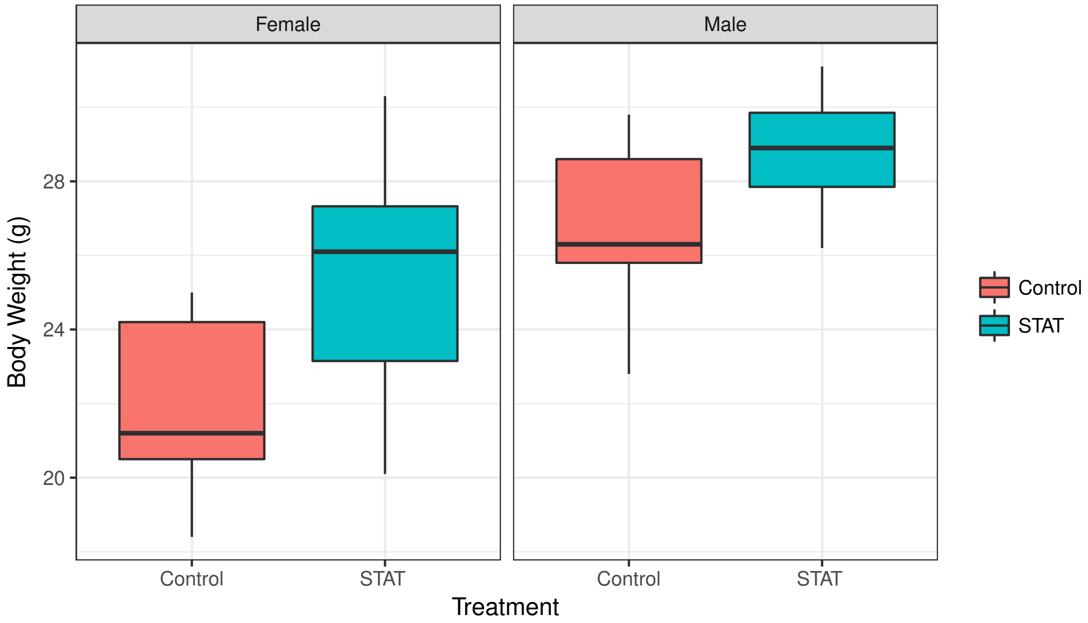


Fig. S4. The box plots of body weights in the subgroups defined by treatment and gender.
